# Matrix linear models for high-throughput chemical genetic screens

**DOI:** 10.1101/468140

**Authors:** Jane W. Liang, Robert J. Nichols, Śaunak Sen

## Abstract

We develop a flexible and computationally efficient approach for analysing high throughput chemical genetic screens. In such screens, a library of genetic mutants is phenotyped in a large number of stresses. The goal is to detect interactions between genes and stresses. Typically, this is achieved by grouping the mutants and stresses into categories, and performing modified t-tests for each combination. This approach does not have a natural extension if mutants or stresses have quantitative or non-overlapping annotations (eg. if conditions have doses, or a mutant falls into more than one category simultaneously). We develop a matrix linear model framework that allows us to model relationships between mutants and conditions in a simple, yet flexible multivariate framework. It encodes both categorical and continuous relationships to enhance detection of associations. To handle large datasets, we develop a fast estimation approach that takes advantage of the structure of matrix linear models. We evaluate our method’s performance in simulations and in an E. coli chemical genetic screen, comparing it with an existing univariate approach based on modified t-tests. We show that matrix linear models perform slightly better than the univariate approach when mutants and conditions are classified in non-overlapping categories, and substantially better when conditions can be ordered in dosage categories. Our approach is much faster computationally and is scalable to larger datasets. It is an attractive alternative to current methods, and provides a natural framework extensible to larger, and more complex chemical genetic screens. A Julia implementation of matrix linear models and the code used for the analysis in this paper can be found at https://bitbucket.org/jwliang/mlm_packages and https://bitbucket.org/jwliang/mlm_gs_supplement, respectively.

## 1 Introduction

High-throughput assays have revolutionalized biology. It was made possible by advances in automation and multiplexing, availability of large and comprehensive collections (such as mutant libraries, and sequenced genomes), and advances in computational and statistical methodology. In this note we consider high-throughput genetic screens which have been deployed for answering complex, large-scale scientific questions. Consider a high throughput genetic screen to observe the fitness of a library of mutants in a variety of growth conditions. Potential goals of such a screen would be to analyze condition × gene interactions or to predict the effect of a new, but related, antibiotic. Matching genes with phenotypes is a particularly valuable application of high-throughput experiments in the age of rapid sequencing technology [van Opijnen and Camilli, 2012]. These techniques can shed light on the physiological roles of partially redundant gene functions [Typas et al., 2011] and the physiological pathways that are involved with responses to different environmental factors [Ivask et al., 2013]. Or, they can reveal relationships between unknown or seemingly unrelated genes [Oh et al., 2011] and provide insights on genes involved in multiple antibiotic resistance [Nichols et al., 2011].

We are now able to run high-throughput genetic screens cost-effectively and in bulk, but the unprecedented scale of these types of studies necessitates the development of generalizable and efficient methods for analyzing their results. Such studies are essentially multivariate problems, but most traditional methods turn them into several univariate problems for computational feasibility. Doing so fails to take advantage of known groupings and correlations in the observations. In the case of the genetic screening example, one can group mutants by gene or gene family: group growth conditions by antibiotic class or temperature; and consider spatial correlation on plates.

We present matrix linear models, which provide a formal statistical framework for encoding such known, but perhaps non-explicit, underlying relationships to enhance detection of associations that might otherwise be masked. This straightforward, multivariate approach can take into account any number of continuous or categorical covariates. Existing methods can encode for different mutant strains and condition types, but are univariate: these approaches are akin to ANOVA or t-tests. In addition to categorical groupings, matrix linear models have the distinct advantage of being able to explicitly model more complex and potentially non-categorical information, such as dosage response levels, null/hotorozygous/homozygous genotypes, and spatial correlation on the plates. (Colonies located on the edge of the plate are expected to exhibit greater growth, since they have fewer neighbors with whom to compete for resources.). Matrix linear models offer flexibility over existing methods analogous to what linear regression offers over t-tests.

Estimation of matrix linear models is fast even in moderately large dimensions. Using simulations and data from an *E. coli* genetic screen [Nichols et al., 2011]. we show that our method produces results comparable to those of the univariate *S* score approach [Collins et al., 2006], but with considerably more efficient computation time. We also analyze the data while encoding for dosage response of growth conditions, to demonstrate the method’s ability to incorporate information from continuous covariates and assess relationships more generally.

This paper is organized as follows. Section 2 introduces a *E. coli* chemical genetic screen data that motivated our method. Section 3 describes the statistical model and estimation. Section 4 evaluates our method using simulated and real datasets. We conclude with a discussion in Section 5.

## 2 *E. coli* Genetic Screening Data

In this high-throughput genetic screening experiment [Nichols et al., 2011] [Shiver et al., 2016]. colony-opacity was recorded for mutant strains grown in high-density on agar plates with a range of conditions. Six plate arrangements of mutants were used, with 1536 colonies grown per plate. The 3983 mutant strains were taken from the Keio single-gene deletion library [Baba et al., 2006]; essential gene hypomorphs (C-terminally tandem-affinity tagged [Butland et al., 2008] or specific alleles); and a small RNA/small protein knockout library [Hobbs et al., 2010]. The colonies were grown in 307 conditions representing different *E. coli* stresses. More than half were antibiotic/ antimicrobial treatments, but they also included other types of conditions, such as temperature and pH. Among the six plate arrangements, a total of 982,902 condition × gene interactions need to be estimated, along with main effects across interactions for the growth conditions and mutant strains. The study aimed to examine the interaction effects between the *E. coli* genes and the growth conditions. This information can be used to study-potential drugs with unknown targets and the mechanism behind drug interactions, as well as identify the genes necessary-to support growth in different conditions [Nichols et al., 2011].

## 3 Model and Estimation

### 3.1 Model

Suppose that *Y* is an *n* × *m* matrix of quantitative colony growth from a high-throughput genetic screen similar to the one described in the previous section. The growth conditions are annotated by *X*_*n*×*p*_ along the *n* rows and the different mutant strains are annotated by *Z*_*m*×*q*_ along the *m* columns. Matrix linear models are thus given by

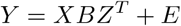

with the main and interaction effects contained in *B*_*p*×*q*_ and errors in *E*_*n*×*m*_. The statistical form of the model model is similar to that used by [Xiong et al., 2011] for genetic analysis of function-valued phenotypes.

If ⊗ denotes the Kronecker product and vec is an operator that stacks columns of a matrix into a single column vector, the vectorized equivalent is

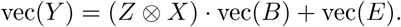

This resembles the familiar linear regression model of *y* = *Xβ* + *ϵ*, and so matrix linear models in this form can theoretically be analyzed using traditional methods for least squares linear regression methods. However, doing so is frequently computationally inefficient or even infeasible.

If the *n* plates, each exhibiting a certain growth condition, are independent with a common covariance matrix, then var(vec(*E*)) = *I_n_* ⊗ Σ. The residual covariance matrix Σ is generally unknown and must be estimated using the data.

### 3.2 Estimation

To obtain least squares estimates, we choose *B* to minimize the residual sum of squares:

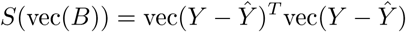

This approach may be viewed as a generalized estimation equations approach [Xiong et al., 2011]. The solution has a closed-form:

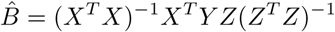

The solution can be viewed as a combination of a least squares on the mutant strains (the *Z* part) and another on the conditions (the *X* part). The resulting estimate is asymptotically unbiased [Liang and Zeger, 1986].

The variance of the estimated coefficient is

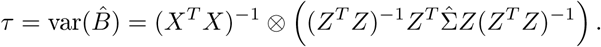

If a consistent estimate of the residual covariance matrix Σ can be obtained, so can a consistent estimate of the variance of the coefficient estimates.

The fitted values are

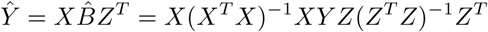

### 3.3 Testing

Similar to the *t*-statistic used to assess the coefficients from the typical (univariate) linear regression model, we can define a test statistic for our method:

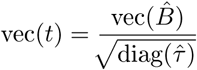

Based on simulations and real data, we empirically observed that the test statistics approximate a skewed *t* distribution. Rather than attempting to specify appropriate parameters for a skewed *t* distribution, we used permutation tests to obtain p-values for our analysis.

Matrix linear models handle only complete data, so all missing values should be dropped, smoothed, or estimated beforehand, as appropriate. The *E. coli* data set had no missing values, so no such considerations were needed. Weighted least squares can be used for heteroscedastic data by weighing plates and/or colonies.

## 4 Simulations Studies and Data Analysis

We applied our method to simulated data and *E. coli* genetic screening data [Nichols et al., 2011]. We compared the results and computation time for matrix linear models and the *S* score, a popular existing method for analyzing high throughput genetic screening data [Collins et al., 2006]. An *S* score is essentially a *t*-statistic comparing the observations for a given mutant and condition with the observations for a given mutant over all conditions. Unlike matrix linear models, which only assume that the rows of *Y* are independent, *S* scores assume that both the rows (plates) and columns (colonies) are independent. So the method is expected to make improvements over *S* scores especially when this assumption is violated, such as when the columns of *Y* (the colonies) are spatially correlated. Matrix linear models go beyond the *S* score ANOVA-like approach and allow for encoding more complex categorical or continuous relationships, similar to linear regression.

### 4.1 Simulation studies

Using the framework of the *X* and *Z* matrices from the *E. coli* data’s six plate arrangements, we simulated data with 1/2 nonzero main effects and 1/4 nonzero interactions drawn from a Normal(0, 2) distribution. The errors were independent and identically distributed from the standard normal distribution. We then applied both our multivariate method and the *S* score’s univariate approach to estimate approximately 180,000 interactions. Using permutation tests, we obtained p-values corresponding to each interaction for each approach. The p-values were then converted into q-values to help account for multiple testing.

To compare the results for each plate arrangement, we plotted the receiver operating characteristic (ROC) curve generated by obtaining true positive rates (TPR) and false positive rates FPR) at varying q-value cutoffs for both methods. The cutoffs were used to determine which q-values corresponded to significant (nonzero) interactions, and these results were then compared to the simulated interactions to obtain the varying TPRs and FPRs. Figure 1 is the ROC plot for the first plate arrangement. The grey-reference line that cuts diagonally from the lower left to the upper right is what we would expect the curve to look like for a method that just produces random noise. Its area under the curve (AUC) is 0.5. A method that performs well will have a curve that closely aligns with the upper left corner and an AUC approaching 1 (its true positive rate will be high even if its false positive rate is low for a given cutoff). See Figure S1 for ROC plots for the remaining five plates.

**Figure 1:**
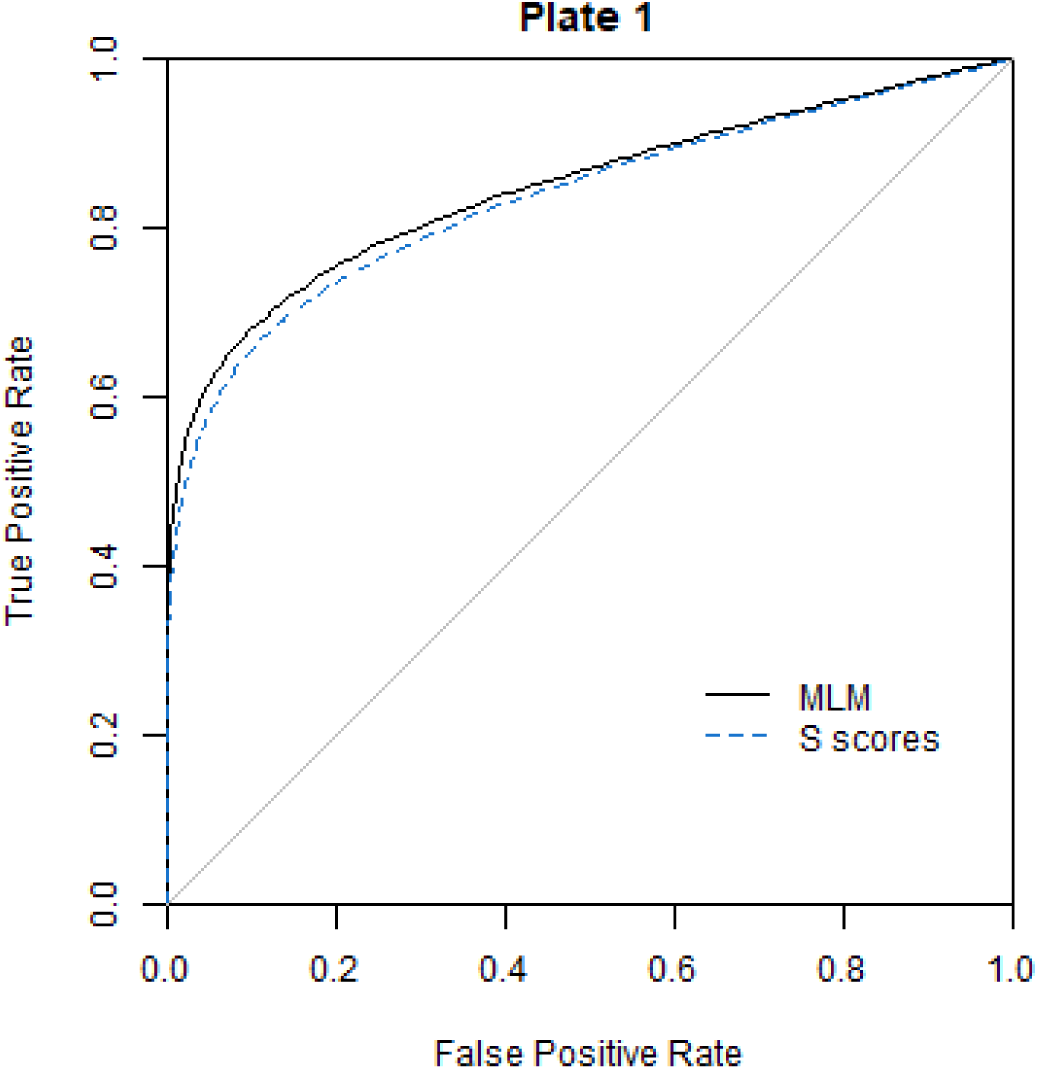
ROC curves comparing MLM to *S* scores applied to data simulated using framework of first plate arrangement. Interactions corresponding to q-values below a given cutoff are considered significant: these results are compared to the simulated interactions. Curves were generated by plotting the TPR and FPR at varying q-value cutoffs. The two methods perform very similarly, with MLM (AUC of 0.845) performing slightly better than the *S* scores (AUC of 0.833).

The AUCs for our method and Collins’s method were 0.845 and 0.833, respectively. Based on both the visual and quantitative summaries, we can observe that matrix linear models perform as least as well as the *S* scores. However, this slight positive difference is consistent across all six plate arrangements [Table 1].

**Table 1:** Area under the curve for simulations based on each of the six plate arrangements. This is computed directly from the ROC curves, as in Figure 1. Matrix linear models outperform Collins’s *S* scores slightly but consistently in each of the six cases.

| Plate | 1 | 2 | 3 | 4 | 5 | 6 |
| --- | --- | --- | --- | --- | --- | --- |
| MLM | 0.845 | 0.843 | 0.853 | 0.852 | 0.847 | 0.853 |
| S scores | 0.833 | 0.838 | 0.852 | 0.850 | 0.840 | 0.852 |

#### 4.1.1 Dosage-response simulation

A more interesting case that illustrates the flexibility and benefits of using matrix linear models is to consider a a genetic screen whose plate conditions have multiple dosage levels. Suppose a given condition has 3 dosage levels. The *S* score approach will analyze these condition × gene interactions separately for each of the dosage levels, e.g. ConditionLevel1 × gene, ConditionLevel2 × gene, and ConditionLevel3 × gene. It is possible to analyze the data analogously using matrix linear models by encoding each of the condition-dosage combinations as separate dummy variables. However, it is also possible for our method to encode this information as dosage response levels for a given condition. Instead of treating this hypothetical condition as essentially three separate conditions with no relationship to each other, we can encode the three dosage response levels together as a single variable corresponding to the condition.

To examine this scenario more closely, we used the *Z* matrix frameworks for mutant strains from each of the *E. coli* data’s six plate arrangements. For each plate, we simulated effects for an experiment with 10 different conditions, each with 3 dosage levels and 3 replicates. 1/4 nonzero interactions were drawn from a Normal(0, 1/2) distribution. For these 1/4 nonzero interactions, we further simulated monotonic dosage effects. We did this by first randomly selecting a direction for the condition’s effect (positive or negative). Then, for the first dosage level, we simulated an effect from an Exponential(0.5) distribution. For the second dosage level, we took the effect from the first dosage level and added that to a random effect drawn from an Exponential(0.5 *α*) distribution, where *α* = 0.8 level, we summed the second dosage level’s effect with a random effect drawn from Exponential(0.5 *α*^2^). The dosage effects for a given condition were then assigned the appropriate direction that was randomly selected in the first step. This simulation can be extended for any number of conditions with any number of monotonic dosage levels.

The ROC curves in Figure 2 illustrate the performance of the dosage-response encoded approach compared with Collins’s *S* scores, which can only encode categorical information, for the first plate arrangement. For each method, the ROC curve was generated by obtaining adaptive Benjamini-Hochberg permutation p-values and varying the cutoff for determining significant interactions. These were then compared to the true simulated interaction effects to calculate the TPR and FPR.

**Figure 2:**
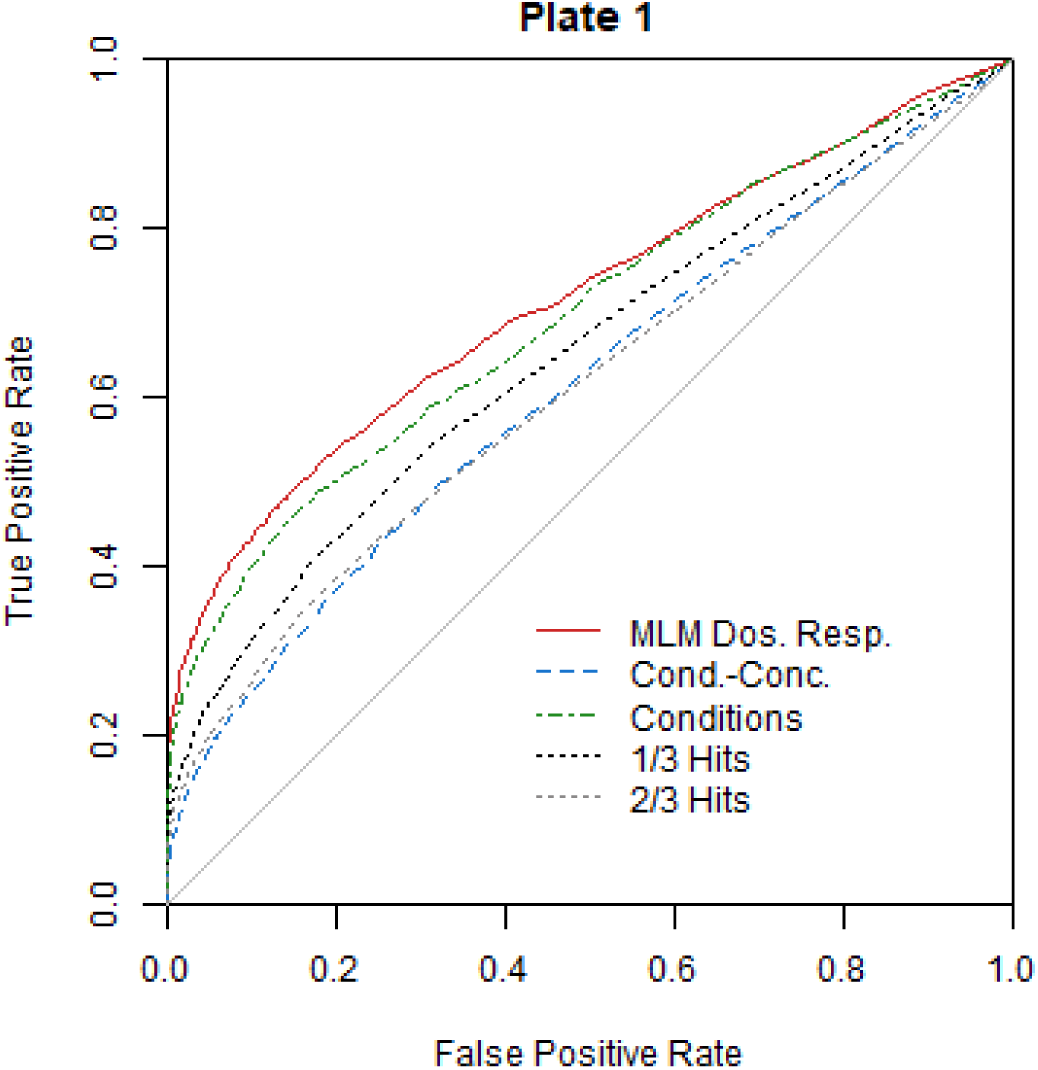
ROC curves for plate 1 simulations comparing dosage-response encoded matrix linear models to categorically encoded *S* scores. Dosage-response encoded matrix linear models outperform all other methods shown. These include *S* scores for data encoded with categorical condition-dosage combinations, for any of the three representations (Cond.-Conc, 1/3 Hits, and 2/3 Hits), as well as *S* scores for data encoded with just the conditions.

- The red solid line plots the results for matrix linear models with dosage-response encoding of the conditions.
- The blue dashed line plots the results for Collins’s *S* scores with categorical encoding of condition-dosage combinations, which is the conventional approach.
- The green dot-dashed line plots the results for Collins’s *S* scores with categorical encoding of only conditions, i.e. all the different dosage levels for a given condition are encoded as one categorical condition.
- The black and grey dotted lines offer an alternate visualization of the Collins’s *S* scores results, encoded for categorical condition-dosage combinations. For a given condition × gene interaction, there are three corresponding *S* scores/ adjusted p-values under this encoding scheme (one for each condition dosage level, which is not information incorporated into the method). When plotting the black dotted “1 /3 Hits” line, a true positive is counted when at least one out of the three adjusted p-values for a significant simulated condition × gene interaction is below the cutoff. Analogously, a false positive is counted when at least one out of the three adjusted p-values for a non-significant simulated condition × gene interaction is below the cutoff. The grey dotted “2/3 Hits” line is generated similarly, but requires at least two out of the three adjusted p-values to be below the cutoff.

The corresponding AUCs for the plate 1 simulation are shown in Table 2.

**Table 2:** AUCs for plate 1 ROC curves plotted in Figure 2. Dosage-response encoded matrix linear models outperform all other methods shown. These include *S* scores for data encoded with categorical condition-dosage combinations, for any of the three representations (Cond.-Conc. 1/3 Hits, and 2/3 Hits), as well as *S* scores for data encoded with just the conditions.

| MLM Dos. Resp. | Cond.-Conc. | Conditions | 1/3 Hits | 2/3 Hits |
| --- | --- | --- | --- | --- |
| 0.714 | 0.612 | 0.694 | 0.650 | 0.612 |

Our proposed dosage-response matrix linear models approach (red solid line) outperforms Collins’s *S* scores encoded with categorical condition-dosage combinations, regardless of representation (blue dashed, black dotted, and grey dotted lines). It also outperforms Collins’s *S* scores when they only encode for the conditions without regard for dosage level (green dot-dashed line). These trends are consistent across simulations based on the other plates’ arrangements [Figure S2 and Table S1]. From an interpretation standpoint, this encoding can be useful if the investigator is interested in the overall effect of a plate condition as opposed to separately considering the different dosage levels. The more general approach to encoding covariates in matrix linear models leads to superior analysis in this situation.

### 4.2 Data analysis

We then applied the our method to each of the six plate arrangements in the *E. coli* genetic screen. The colony opacities were standardized by subtracting the median colony opacity of each plate (which has multiple mutant strains growing under a given set of conditions) and dividing by the IQR.

#### 4.2.1 Computational considerations

When we ran our matrix linear model and *S* score implementations (encoded with condition-dosage combinations) on the entire data set of six plates, the latter required significantly more computation time [Tables 3 and 4]. A computer with 128 GB memory and a 3.00 GHz dual-core processor was used to obtain the times as averages of 10 runs. Matrix linear models only take about three and a half seconds to estimate the roughly 1 million interactions, plus main effects [Table 3]. In comparison, Collins’s *S* scores require around two and a half minutes. Two and a half minutes is still fairly reasonable; however, much greater computation time is needed to obtain permutation p-values. If the p-values are calculated based on 1000 permutations (parallelized over 5 cores), matrix linear models take just over eighteen minutes [Table 4]. *S* scores require over eight hours to complete the same procedure (again, parallelized over 5 cores). This dramatic difference in computation time can have a considerable impact on the scope and feasibility of analyzing such data sets.

**Table 3:** Computation time to estimate condition × gene interactions, plus main effects, for each plate (a total of about 1 million interactions). Matrix linear models are considerably less computationally expensive.

| Plate | 1 | 2 | 3 | 4 | 5 | 6 | Total |
| --- | --- | --- | --- | --- | --- | --- | --- |
| MLM | 0.52 | 0.47 | 0.67 | 0.61 | 0.52 | 0.50 | 3.29 sec |
| *S* scores | 22.81 | 20.83 | 30.52 | 29.94 | 23.82 | 24.08 | 151.99 sec |

**Table 4:** Computation time to estimate permutation p-values for the condition × gene interactions for each plate (a total of about 1 million interactions). The computational advantages of matrix linear models are even more apparent when permutation p-values are desired.

| Plate | 1 | 2 | 3 | 4 | 5 | 6 | Total |
| --- | --- | --- | --- | --- | --- | --- | --- |
| MLM | 2.82 | 2.78 | 3.46 | 3.28 | 2.82 | 3.00 | 18.16 min |
| *S* scores | 77.62 | 72.59 | 93.37 | 94.22 | 76.31 | 77.32 | 491.42 min |

#### 4.2.2 Auxotroph Analysis

To assess whether our method identifies significant interactions in the expected manner, we analyzed auxotrophs. Auxotrophs are mutant strains that have lost the ability to synthesize a particular nutrient required for growth. These might include knockout strains for a certain amino acid. Since they should experience little to no colony growth under specific conditions where the required nutrient is not present, we expect negative interactions between auxotrophic mutants and minimal media growth conditions. Auxotrophs are useful as controls, since the phenotype under particular conditions for a mutant strain is typically not known.

In the original univariate analysis of the colony size data. Nichols et al., empirically identified 102 auxotrophs [Nichols et al., 2011]. Likewise, a previous study of the Keio Collection auxotrophs, based on colony size, found 238 auxotrophs, 110 of which were mutants included in this data set [Joyce et al., 2006]. Nichols et al., and Joyce et al., found a 70% overlap, despite significant experimental differences (e.g. growth in liquid vs. solid media).

In a similar fashion, we empirically identified auxotrophs based on the matrix linear model estimates. We did this by obtaining the quantiles of the MLM interaction scores for each mutant strain under minimal media conditions. Mutants whose 95% quantile for interaction scores with minimal media conditions fell below zero were classified as auxotrophs. Our auxotrophs had a 83.33% overlap with the Nichols et al., auxotrophs and a 71.82% overlap with Joyce et al., The slightly larger intersection of auxotrophs when comparing with Nichols et al., vs comparing with Joyce et al., is to be expected, since we are analyzing the same data set. As noted above, Joyce et al., was a separate study with experimental differences. While not all of the matrix linear model interactions between the Nichols et al., and Joyce et al., auxotrophs and minimal media conditions were negative, the vast majority of them were. Figures 3, S4 are visualizations of the distributions of each auxotroph’s interaction scores across minimal media conditions. The interaction scores and plotted as points, and the median for each auxotroph is plotted as a horizontal bar. Most fall below zero, which is what we would expect.

**Figure 3:**
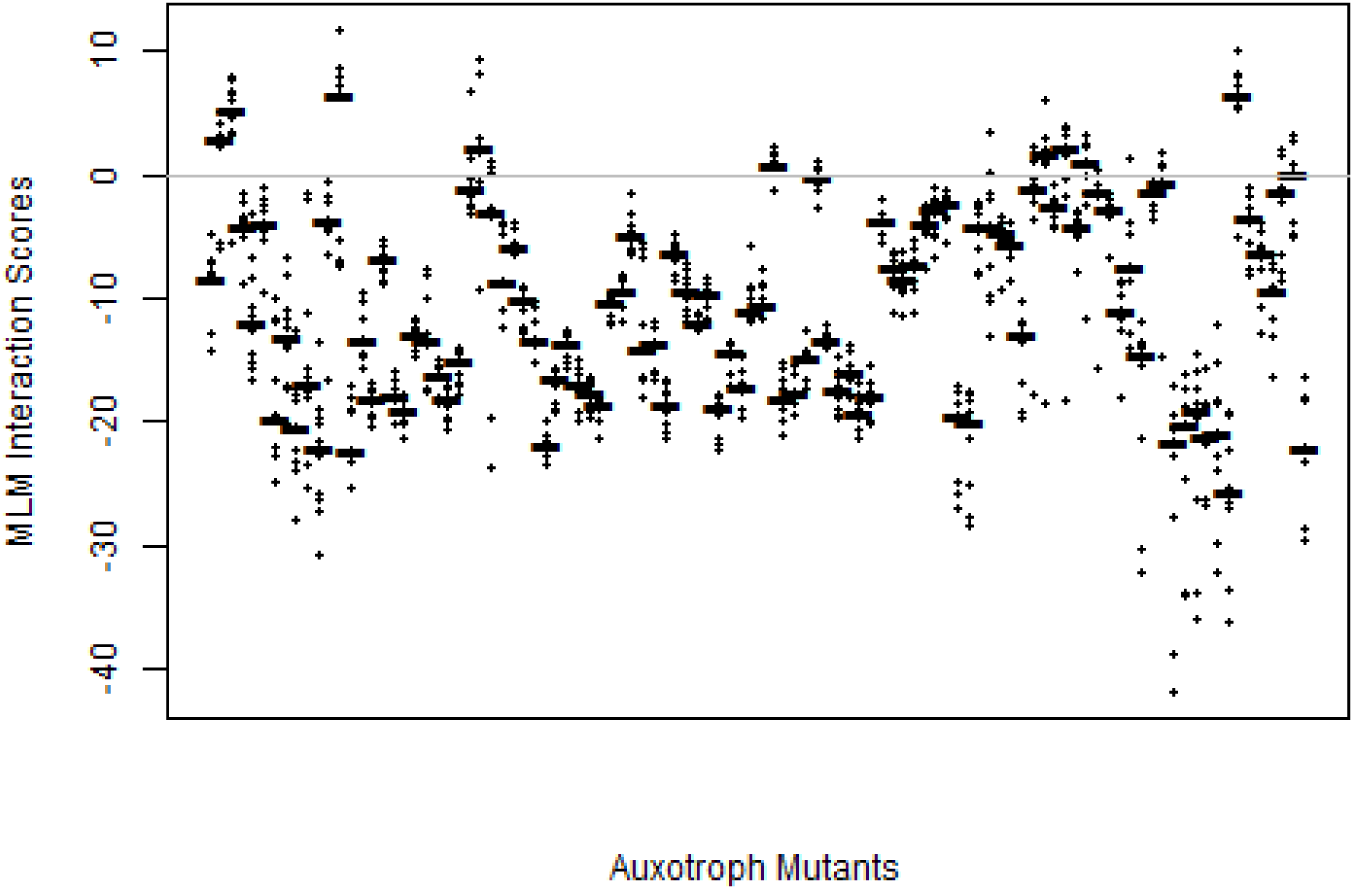
Distributions of matrix linear model interaction estimates for auxotrophs identified by [Nichols et al., 2011] over minimal media conditions. The Nichols et al., auxotrophs are plotted along the horizontal axis. The MLM interactions between the auxotrophs and minimal media conditions are plotted along the vertical axis, with the horizontal bars indicating the median value. Most interactions fall below zero, indicating little growth.

However, some of the discrepancy may be due to differences between analyzing colony opacity, as we did, and analyzing colony size, as Nichols et al., and Joyce et al., did. Kritikos et al., ran a study of the Keio collection, but analyzed colony opacity and made their raw *S* scores publicly available [Kritikos et al., 2017]. We used the Kritikos et al., *S* scores to replicate the auxotroph-identifying process for MLM t-statistics. Of the nineteen Nichols et al., auxotrophs that MLM was not able to identify, the Kritikos et al., *S* scores were also unable to detect twelve. This result suggests that about two-thirds of the Nichols et al., auxotrophs that MLM was unable to detect can be accounted for by differences in the types of measurements used to quantify growth.

Of the remaining five auxotrophs that MLM failed to find but that the Kritikos et al., *S* scores identified, fepC is a mutant that results in the loss of ferric enterobactin uptake. There are minimal media conditions for both “high iron” and “low iron”. The MLM t-statistics are generally positive for interactions between fepC and both iron growth conditions; the Kritikos et al., *S* scores are positive only for “high iron”. Thus. fepC might be considered a borderline case in auxotroph determination depending on whether or not it exhibits growth under the “low iron” condition. Additionally, there were two auxotrophic mutants that were found through analysis of the MLM t-statistcs, but not through analysis of the Kritikos et al., *S* scores.

22 out of the 31 Joyce et al., auxotrophs that the MLM t-statistics were unable to find were also undetectable to the Kritikos et al., *S* scores. Once again, this suggests that about two-thirds of the discrepant auxotrophs can likely be explained by differences between analyzing colony size and colony-opacity.

[Kritikos et al., 2017]

As an alternative visualization, consider ROC plots [Figure 4, S5] that assess the ability of matrix linear models to correctly identify auxotrophs found by Nichols et al., and Joyce et al., To get the TPR and FPR for Figure 4, we took the auxotrophs identified by Nichols et al., to be the “true” auxotrophs. We then obtained TPRs and FPRs by varying cutoffs for the median minimal media interaction score for the auxotrophs that we identified. Figure S5 was obtained analogously for the Joyce et al., auxotrophs. The AUCs were 0.884 and 0.824, respectively; there is high concordance between the three sets of auxotrophs.

**Figure 4:**
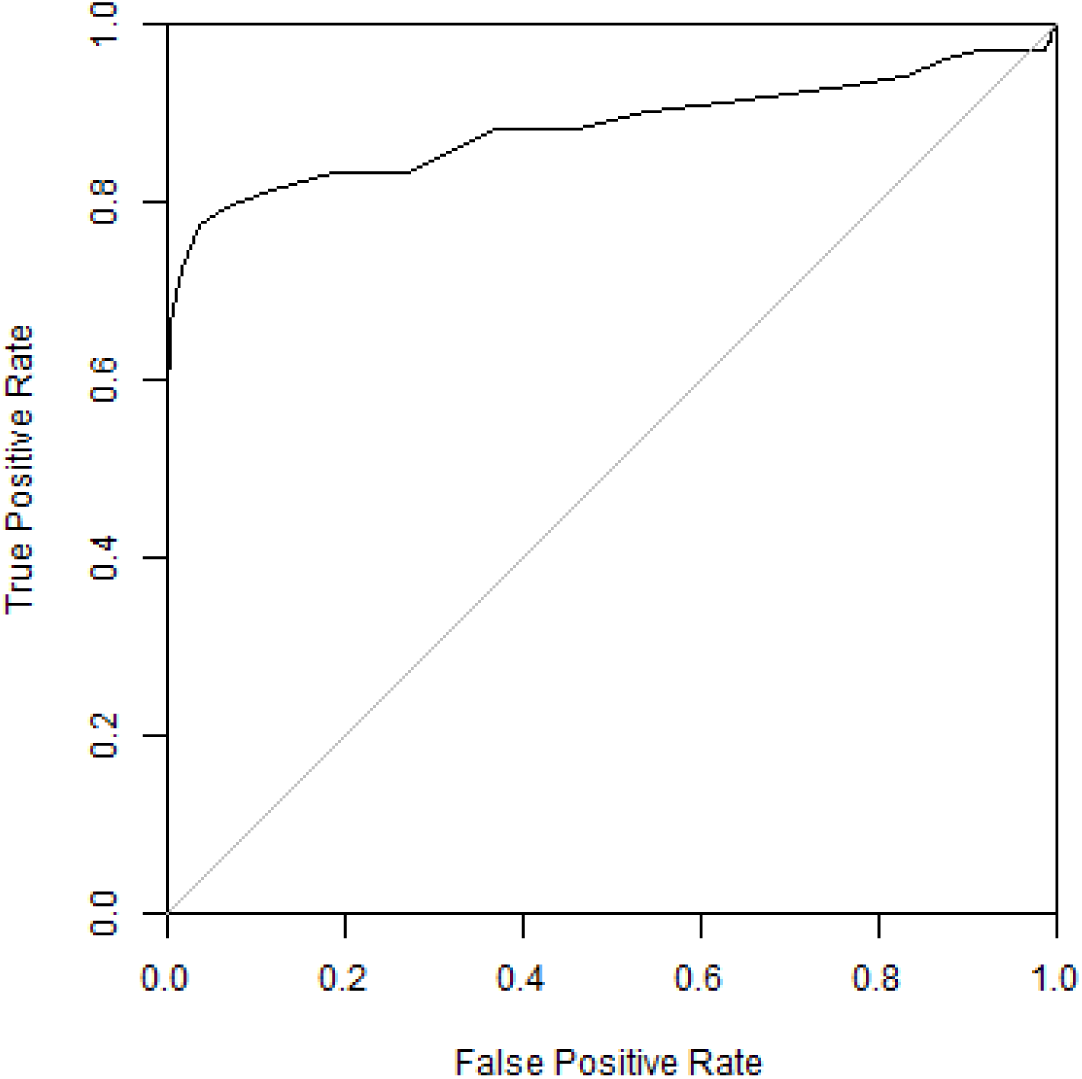
ROC curve for the auxotrophs we empirically identified, compared against those identified by [Nichols et al., 2011] as the reference. TPRs and FPRs were calculated based on the median minimal media interaction score for the each of the auxotrophs we identified, at varying cutoffs. The AUC was 0.884.

### 4.3 Dosage-response analysis

We also analyzed the data set by running matrix linear models with dosage-response levels encoded in the *X* matrix growth conditions. It should be noted that many of the conditions in the Nichols et al., data set had only one dosage level, which makes it somewhat less-than-ideal for illustrating this encoding approach. We then compared the dosage-response results with the results from applying matrix linear models and Collins’s *S* scores on data conventionally encoded for condition-dosage combinations.

Figure 5 examines the performance of these three approaches for analyzing the first plate arrangement. (The other five plate arrangements, shown in Figure S3, produced similar results.) For each of the three methods, we used permutation tests to obtain adaptive Benjamini-Hochberg adjusted p-values corresponding to each interaction. We then plotted the proportion of adjusted p-values below varying thresholds to generate the three curves. The dosage-response-encoded matrix linear models were able to detect more significant interactions (adjusted p-values below a given cutoff) than Collins’s method at nearly every threshold. For lower cutoffs, this continuous encoding strategy also detects more significant interactions than matrix linear models with categorical encoding.

**Figure 5:**
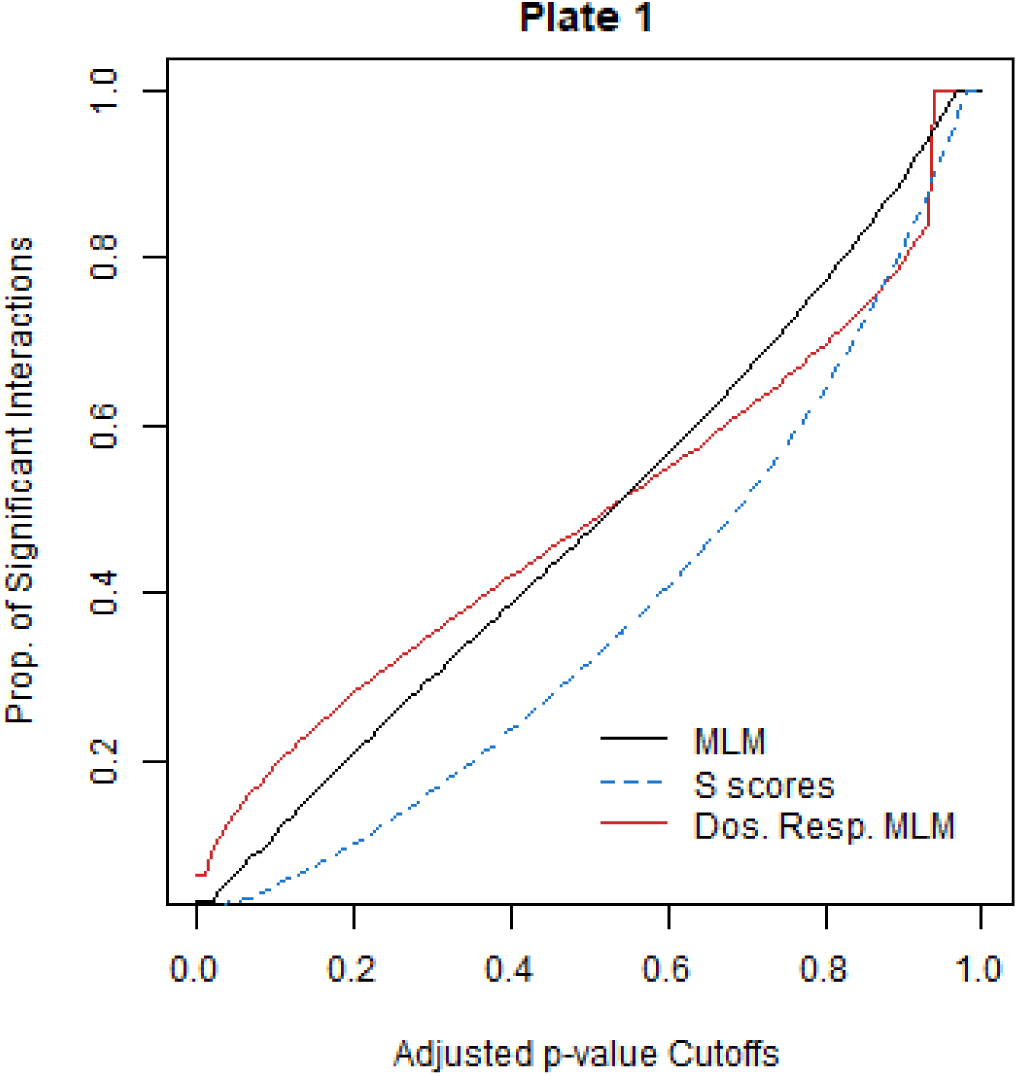
Comparing the proportion of significant interactions detected when encoding for dosage response and when encoding dosage-condition combinations categorically, for the first plate arrangement. The former is only possible in matrix linear models. The latter is shown for both matrix linear models and *S* scores. The curves were generated by obtaining the adaptive Benjamini-Hochberg adjusted p-values from permutation tests for each method and identifying the proportion of adjusted p-values below varying cutoffs.

This simple example illustrates the potential gains in performance and detection of significant interactions when taking advantage of matrix linear models’s regression-like ability to encode for complex and continuous covariates. Encoding the conditions in this manner also provides interpretable results about the effects of dosage in the interactions as well as the effects of the conditions themselves.

## 5 Discussion

We have presented matrix linear models, a simple framework for encoding relationships and groupings in high-throughput genetic screenings. This approach is computationally efficient, requiring significantly less time to run than the univariate *S* score method [Collins et al., 2006]. The speed advantage is especially noticeable when performing analysis on larger data sets or doing repeated runs, as for a permutation test.

Matrix linear models can also improve the detection of interaction effects that might otherwise be masked. By evaluating our method alongside the *S* score approach when applied to simulations and an *E. coli* genetic screen [Nichols et al., 2011]. we show that we achieve comparable results at much less computational expense. Furthermore, unlike the *S* score, matrix linear models are not limited to encoding categorical groupings. In this way, the relationship between *S* scores and matrix linear models can be thought of as being analogous to that between ANOVA/t-tests and linear regression. Analysis of the simulated and *E. coli* data when encoding for multi-level condition dosages demonstrates how matrix linear models can provide a more flexible and powerful approach that can analyze scenarios with non-categorical covariates.

In this paper, we discussed a generalized estimating equation approach to estimation that uses least squares as the computational engine. Several extensions are possible. We may want a robust estimation procedure that downweights extreme observations (least squares is well-known to be sensitive to outliers). This can be achieved by modifying the loss function from a sum of squares to a robustified version (such as Huber’s loss function)[Hastie et al., 2009]. The resulting optimization problem is expected to be more complex, however. Another extension might be to fit penalized matrix linear models with a L_1_ penalty. This optimization problem is also challenging, and we expect to report progress in future work. Finally, although we were motivated by high-throughput chemical genetic screens, many other high throughput data such as metabolomic data or cancer cell line drug screening have a similar structure and might benefit from a similar approach.

A Julia implementation of matrix linear models can be found at https://bitbucket.org/jwliang/mlm_packages. The code to perform the analysis and generate the figures in this paper is available at https://bitbucket.org/jwliang/mlm_gs_supplement.

## 6 Acknowledgements

We thank Dr. Carol Gross of UCSF for allowing us to use and share the chemical genetic screen data. We thank Anthony Shiver of Stanford University for help accessing, processing, and interpreting the genetic screen data. This work was started when JWL was a summer intern at UCSF. and continued when she was a scientific programmer at UTHSC. We thank both UCSF and UTHSC for funding, and a supportive environment for this work. SS was partly supported by NIH grants GM070683. GM078338. GM123489. and DA044223.

## Supplemental figures and tables

**Table S1:** AUCs for dosage-response simulation based on plates 2-5. Dosage-response encoded matrix linear models outperform all other methods shown within each plate. These include *S* scores for data encoded with categorical condition-dosage combinations, for any of the three representations (Cond.-Conc, 1/3 Hits, and 2/3 Hits), as well as *S* scores for data encoded with just the conditions.

| Plate | 2 | 3 | 4 | 5 | 6 |
| --- | --- | --- | --- | --- | --- |
| MLM Dos. Resp. | 0.707 | 0.779 | 0.688 | 0.736 | 0.720 |
| Cond.-Conc. | 0.623 | 0.663 | 0.591 | 0.644 | 0.632 |
| Conditions | 0.683 | 0.753 | 0.666 | 0.713 | 0.705 |
| 1/3 Hits | 0.657 | 0.713 | 0.625 | 0.673 | 0.683 |
| 2/3 Hits | 0.634 | 0.666 | 0.588 | 0.655 | 0.630 |

**Figure S1:**
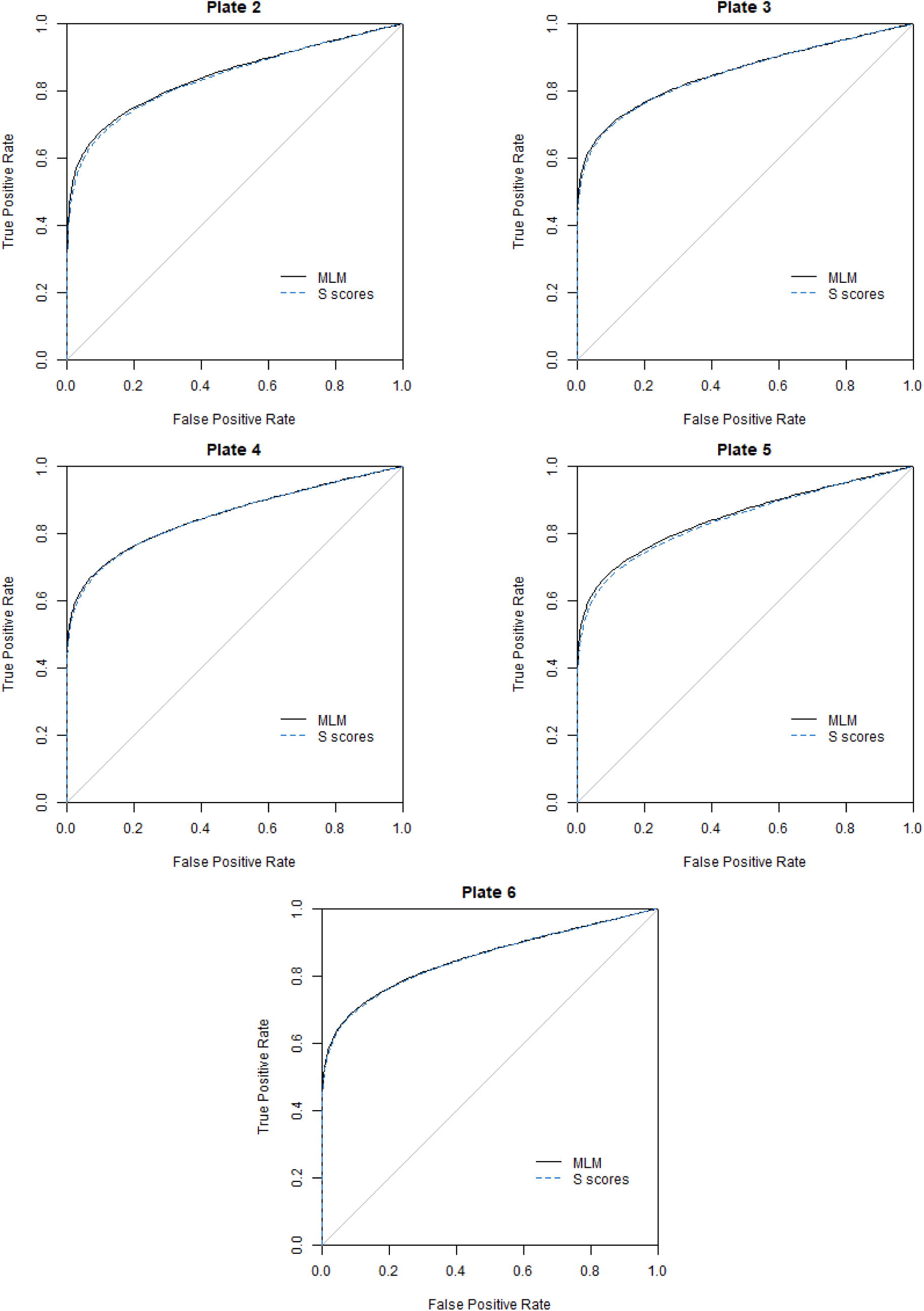
ROC curves comparing MLM to *S* scores applied to data simulated using frameworks of plates 2-5. Interactions corresponding to q-values below a given cutoff are considered significant: these results are compared to the simulated interactions. Curves were generated by plotting the TPR and FPR at varying q-value cutoffs

**Figure S2:**
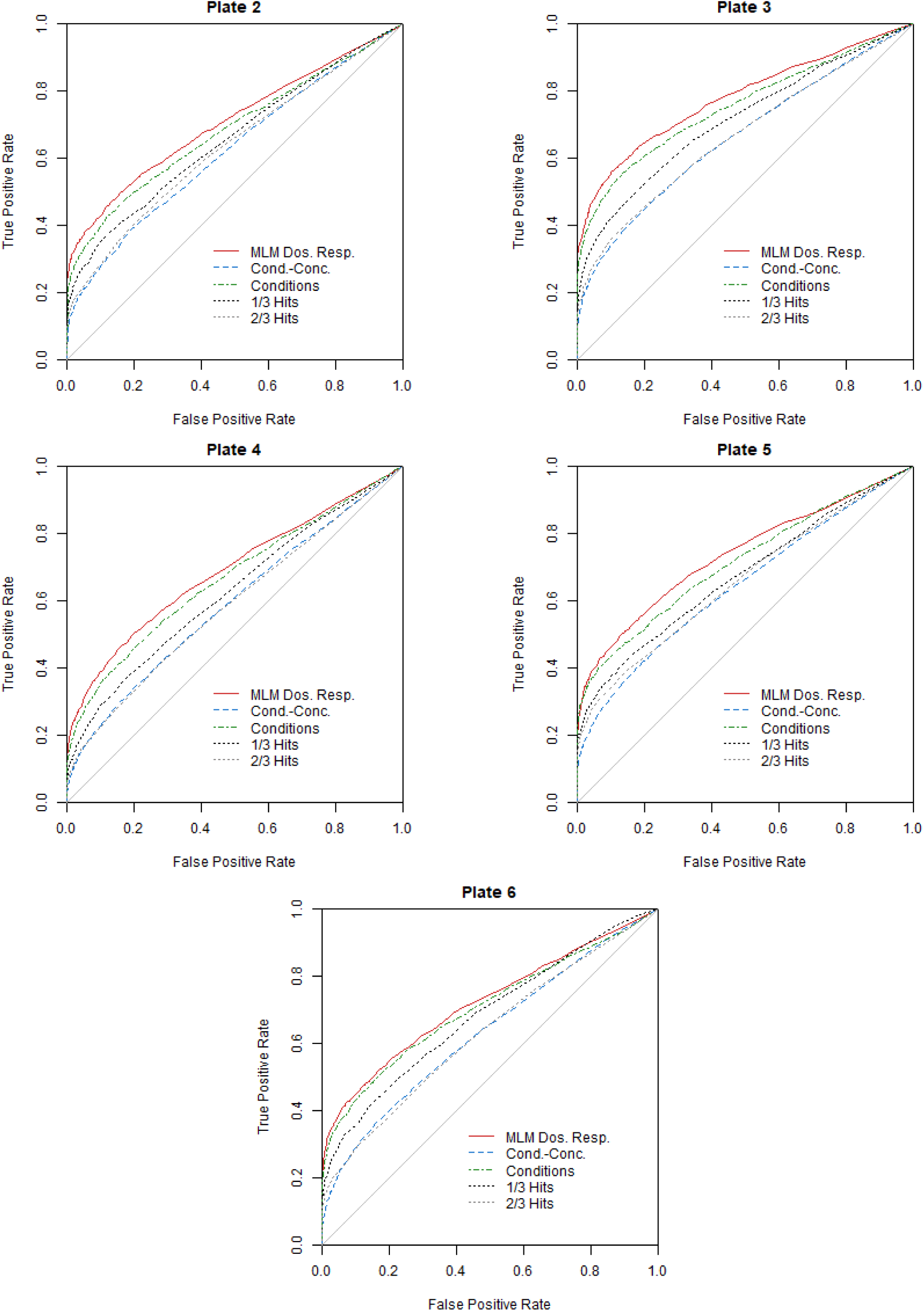
ROC curves for plate 2-5 simulations comparing dosage-response encoded matrix linear models to categorically encoded *S* scores. Dosage-response encoded matrix linear models outperform all other methods shown. These include *S* scores for data encoded with categorical condition-dosage combinations, for any of the three representations (Cond.-Conc, 1/3 Hits, and 2/3 Hits), as well as *S* scores for data encoded with just the conditions.

**Figure S3:**
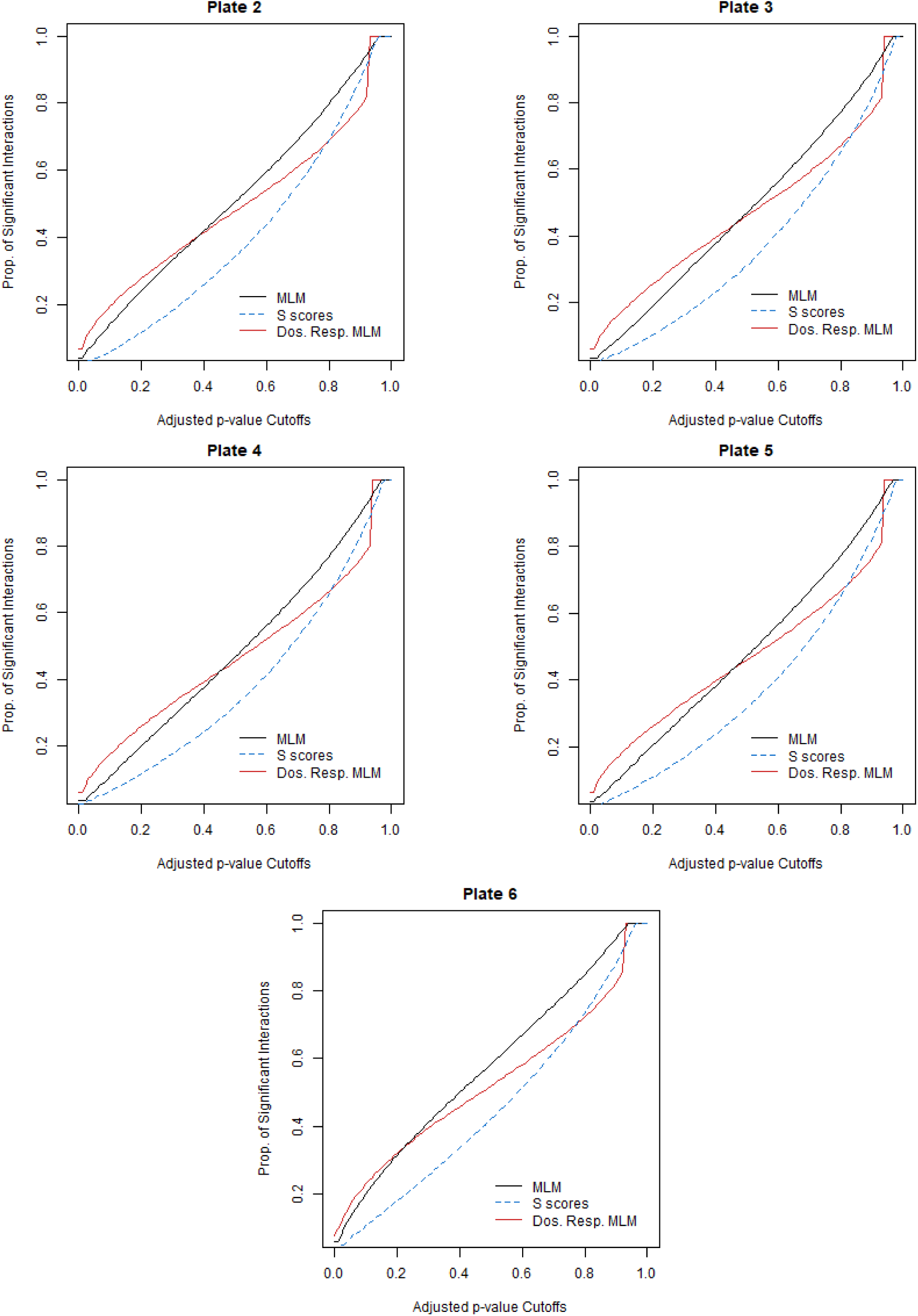
Comparing the proportion of significant interactions detected when encoding for dosage response and when encoding dosage-condition combinations categorically, for plates 2-5. The former is only possible in matrix linear models. The latter is shown for both matrix linear models and *S* scores. The curves were generated by obtaining the adaptive Benjamini-Hochberg adjusted p-values from permutation tests for each method and identifying the proportion of adjusted p-values below varying cutoffs.

**Figure S4:**
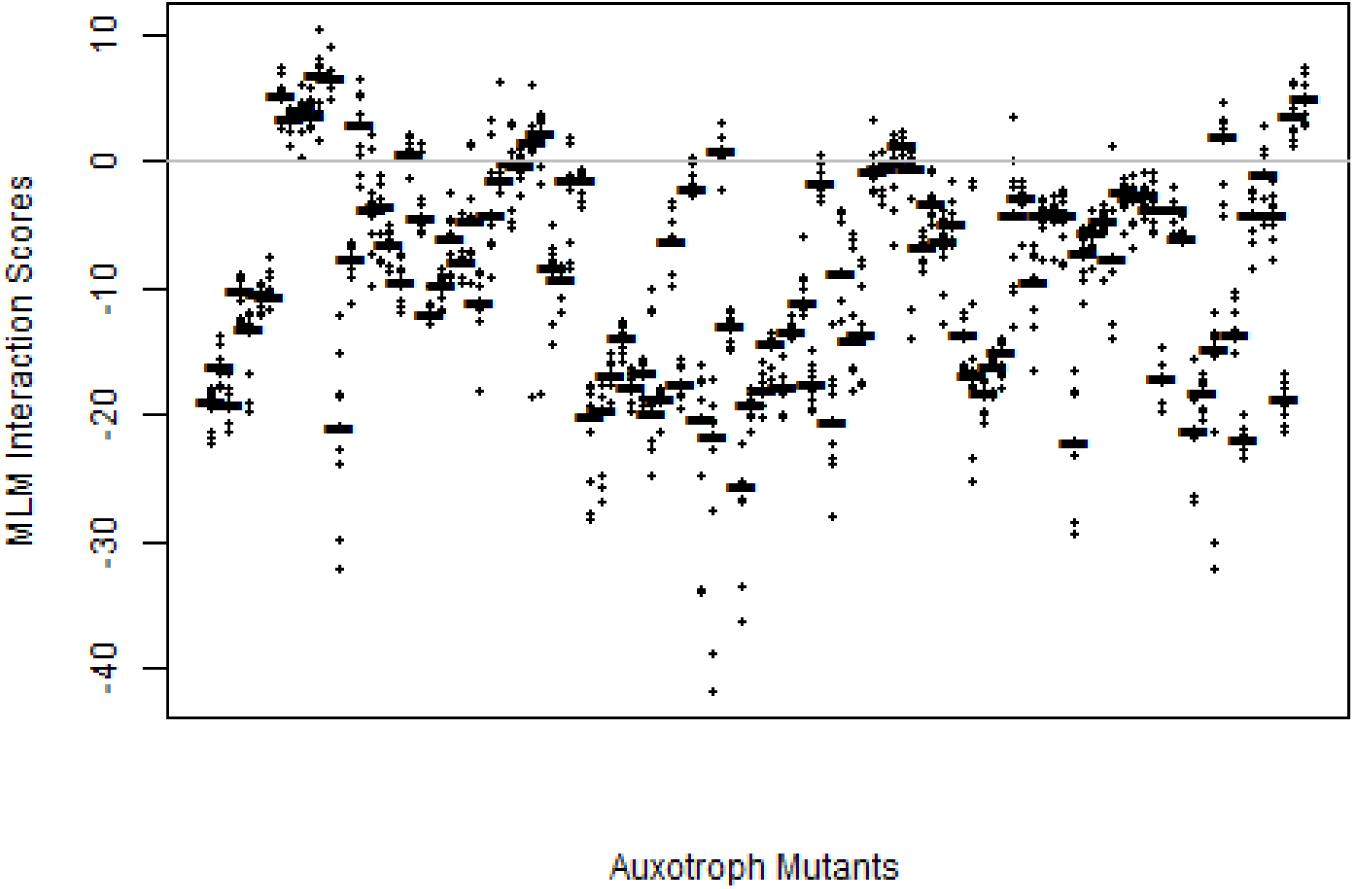
Distributions of matrix linear model interaction estimates for auxotrophs identified by [Joyce et al., 2006] over minimal media conditions. The Joyce et al., auxotrophs are plotted along the horizontal axis. The MLM interactions between the auxotrophs and minimal media conditions are plotted along the vertical axis, with the horizontal bars indicating the median value. Most interactions fall below zero, indicating little growth.

**Figure S5:**
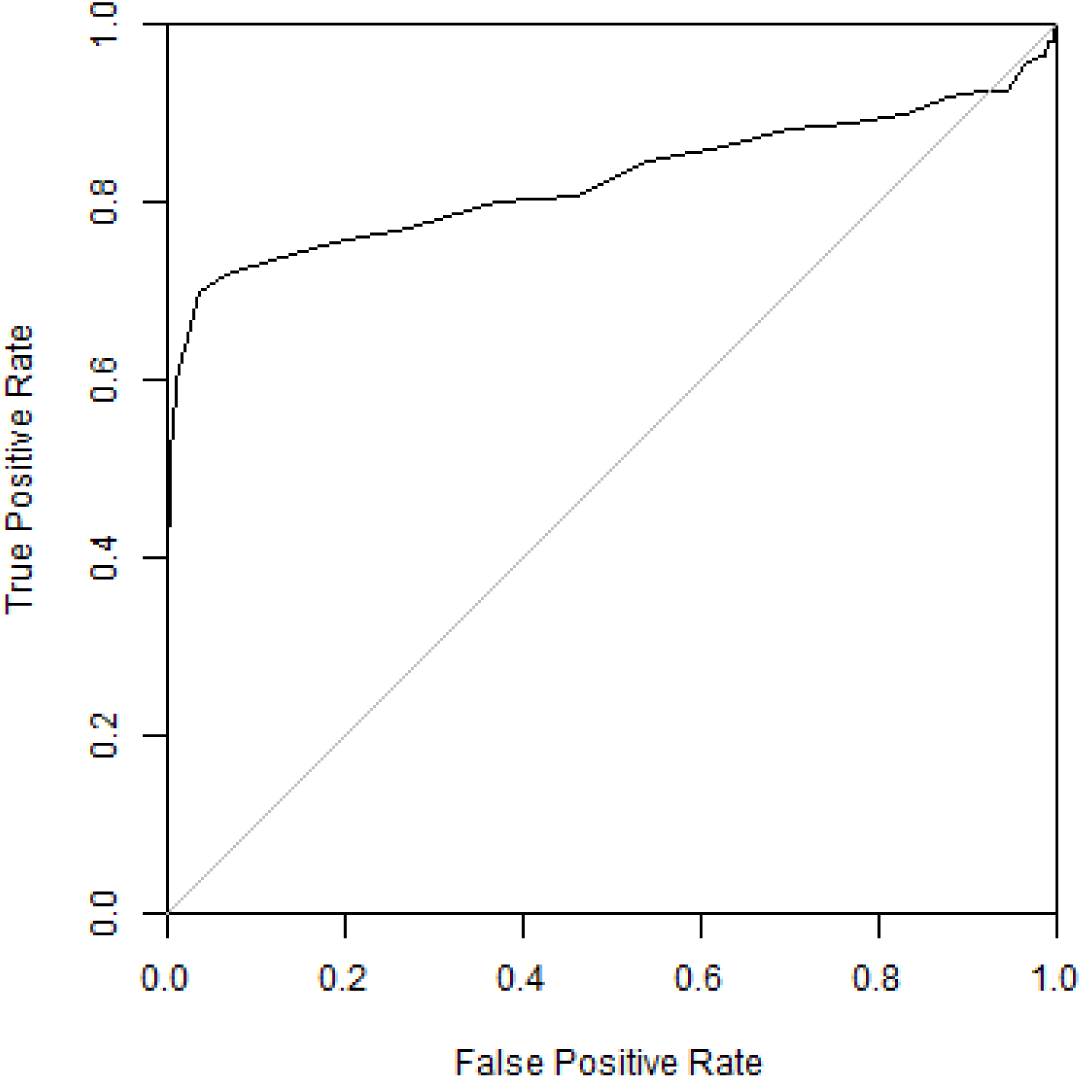
ROC curve for the auxotrophs we empirically identified, compared against those identified by [Joyce et al., 2006] as the reference. TPRs and FPRs were calculated based on the median minimal media interaction score for the each of the auxotrophs we identified, at varying cutoffs. The AUC was 0.824.

